# Microbial Differences Accurately Identifies Global SERT KO Phenotype in Mice

**DOI:** 10.1101/2024.11.08.622711

**Authors:** Madelaine Leitman, Will Katza, David Zhang, Shrey Pawar, Simer Shera, Laura Hernandez, Tien S. Dong

## Abstract

Altered serotonin signaling is a well-established contributor to depression, with the serotonin transporter gene (SERT) playing a critical role in regulating serotonin reuptake. Mice lacking SERT (SERT -/-) serve as a robust model for depression, exhibiting significant depressive-like behaviors compared to littermate wild-type (SERT +/+) controls. In this study, we aimed to determine the relationship between gut microbiota composition and depressive behaviors in SERT -/- mice. Behavioral assays, including the Forced Swim Test (FST) and Tail Suspension Test (TST), confirmed that SERT -/- mice exhibited significantly increased immobility times compared to SERT +/+ mice (FST: p = 0.004; TST: p = 0.080), consistent with a depressive phenotype. Utilizing littermate controls, shotgun metagenomic sequencing of fecal samples revealed significant differences in alpha diversity between the two groups of mice, as measured by the Shannon entropy index (p = 0.05). Additionally, our bacterial co-occurrence network analysis uncovered distinct structural differences in microbial interactions between SERT -/- and SERT +/+ mice (p = 0.001), suggesting shifts in microbiome stability and functionality between the groups. We created a microbial depression score utilizing the top five bacteria taxa that were differentially abundant between SERT -/- and SERT +/+ mice: *Clostridium sp. MD294, Acetatifactor MGBC165152, Desulfovibrio MGBC129232, Oscillibacter MGBC161747, and Schaedlerella MGBC000001.* This microbial depression score correlated strongly with immobility times in the FST (r = 0.705, p < 0.0006) and TST (r = 0.401, p < 0.09). A random forest classifier based on these taxa accurately distinguished SERT -/- from SERT +/+ mice (accuracy = 0.82). These findings suggest that gut microbial species composition is highly associated with depressive-like behaviors in SERT -/- mice, likely via alterations in serotonin signaling pathways, and may offer potential targets for microbiome-based interventions in depression.

## Introduction

Serotonin, also known as 5-hydroxytryptamine (5-HT), is a key neurotransmitter implicated in a wide range of neuropsychiatric disorders, including depression. Serotonin synthesis begins with the essential amino acid tryptophan, the sole precursor for serotonin production both in the raphe nuclei of the brain and the enterochromaffin cells of the intestinal mucosa, the primary sites of central and peripheral serotonin production, respectively ^1^. Although serotonin’s role in the central nervous system (CNS) is well-recognized for its regulation of mood, sleep, and appetite, approximately 95% of the body’s serotonin is produced peripherally in the gastrointestinal tract ^1–3^. Once exerting its effects, serotonin is cleared from the synaptic cleft by the transmembrane serotonin transporter (SERT), a protein encoded by the SLC6A4 (SERT) gene. Terminating serotonin’s signal by removing it from the synaptic cleft prevents continuous receptor activation, which is crucial for mood regulation and overall emotional balance. Dysfunction in SERT leads to elevated extracellular serotonin levels, which paradoxically lead to downregulation of serotonin receptors over time and desensitization of serotonergic pathways^4^. This is a compensatory mechanism to counteract excessive serotonin presence, reducing the sensitivity to serotonin in order to prevent overstimulation of the receptors.

Research has demonstrated that polymorphisms in the SERT gene are strongly associated with depressive phenotypes in both humans and mice^5^. Humans that are homozygous for a short (14 repeats) SERT allele have decreased levels of SERT and are more susceptible to depression relative to those with the long (16 repeats) SERT allele ^6^. Similarly, the SERT knockout (SERT -/-) mouse model has an absence of functional SERT, resulting in persistent serotonin elevation and behavioral phenotypes that closely mimic human depression. These mice behavioral phenotypes include behavioral despair, anhedonia, exaggerated stress responses, and social withdrawal ^7–10^. By affecting serotonin reuptake, the SERT -/- model is a crucial tool in understanding the mechanisms underlying depression and in the development of targeted antidepressant therapies.

Emerging evidence suggests that the gut microbiota influence brain function and mood regulation through interactions with multiple physiological systems, collectively known as the brain-gut-microbiota axis ^11–13^. This axis involves bidirectional communication between the gut microbiota and the enteric nervous system (ENS), central nervous system (CNS), immune system, and endocrine system, with serotonin acting as a key mediator ^11^. Particularly, the gut microbiome plays a critical role in regulating serotonin levels, which, in turn, also shape the composition and activity of the gut microbiota. Beyond their well-known roles in digesting complex carbohydrates, shaping the immune system, and preserving the integrity of the intestinal barrier, gut microbes manipulate host serotonergic pathways through interactions with enterochromaffin cells ^14,15^. In fact, accumulating studies have shown changes in the microbial communities of depression patients compared to healthy individuals. Although there are conflicting results, recurrent patterns include an enrichment of pro-inflammatory bacteria and a depletion of anti-inflammatory bacteria, supporting the inflammatory hypothesis of depression ^16^. While it is known that gut microbiota can affect serotonin pathways, the exact species and mechanisms by which these changes affect SERT -/- mice’s depressive behaviors remain unclear ^17,18^. We aim to explore gut microbial differences between SERT -/- and littermate wild-type (SERT +/+) mice and how these changes correlate with behavioral despair. By identifying microbial species linked to disrupted serotonin signaling and depressive behavior, this study could pave the way for microbiome-targeted therapies in treating depression.

## Methods

### Mice

SERT -/- and SERT +/+ mice of C57BL/6 background (purchased from Jackson Laboratory) were bred in-house at UCLA. Animals were housed in autoclaved polypropylene cages with corncob bedding under a specific pathogen-free environment under a 12 h light/dark cycle. Genotyping was carried out after tail clipping on 4 weeks old mice. SERT -/- mice were homozygous knock outs of the SLC6A4 gene. The mice were housed by litter, sex and genotype until 11–12 weeks of age when fecal samples were collected. All animal studies were performed in accordance with institutional guidelines and regulations.

### Forced Swim Test

To measure depressive behavior, mice underwent the Forced Swim Test (FST) (Figure 1A). The tanks were filled with tap water at 23-25°C, checked with an infrared thermometer, and the test began once the water reached the desired level. White noise was generated to mask external sounds, maintaining consistent conditions for all animals. Mice were gently placed in the water, and testing lasted six minutes, though only the final four minutes were analyzed, as mice are typically more active during the initial phase. For behavioral scoring, video footage of the test was uploaded for analysis. Mobility and immobility times were calculated, with immobility defined as any behavior other than movements necessary to keep the head above water. An on-screen stopwatch was used to record movement times, with the observer blinded to group identities to prevent bias. Statistical analysis was completed in R (R version 4.4.1) ^19^, where the Mann-Whitney U test was applied using the *wilcox.test* function. A threshold of 0.05 was applied to determine significance. Visualizations were produced using *ggplot2* in R.

**Figure 1:**
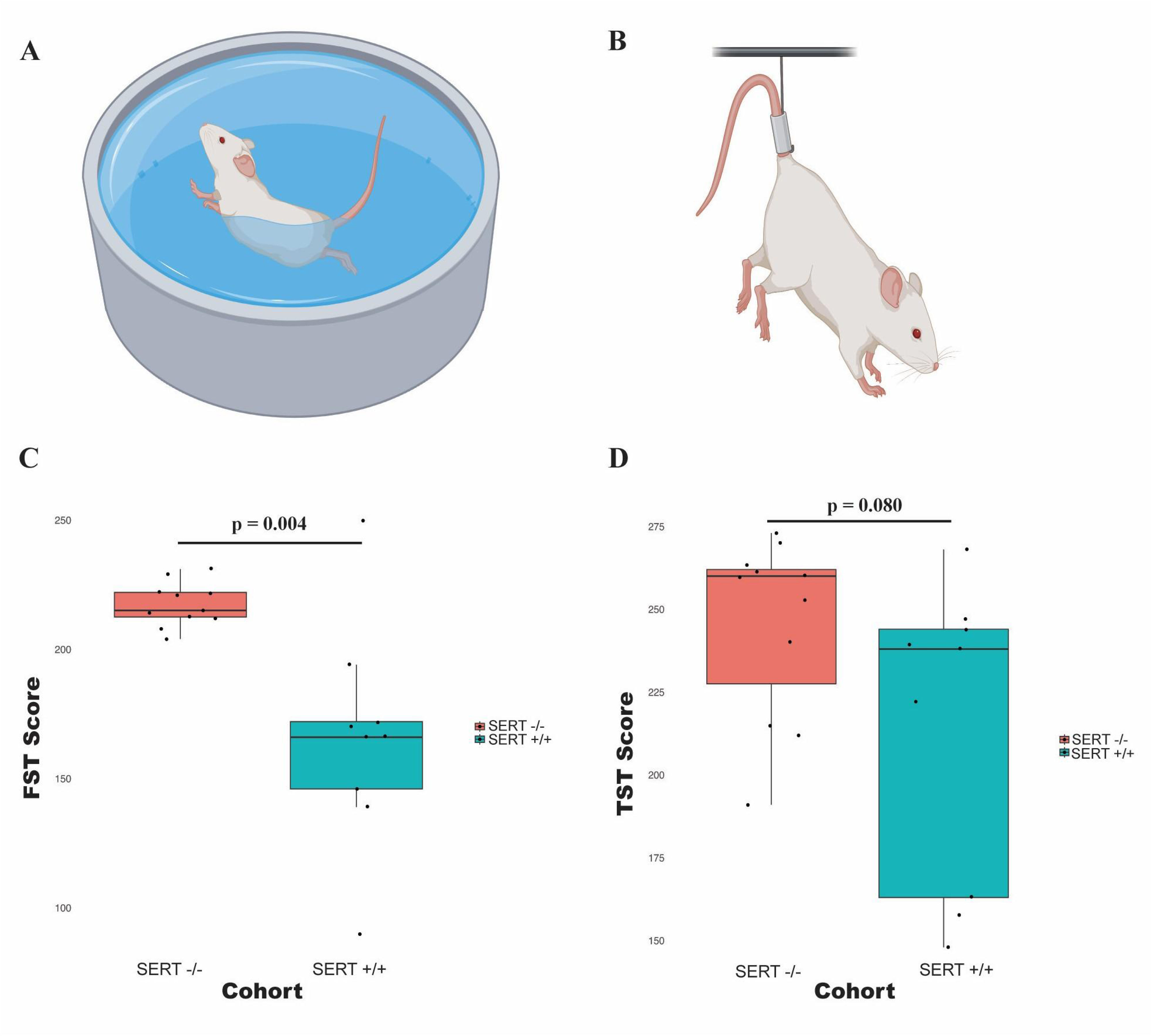
Behavioral assessments of depressive-like phenotypes in SERT -/- and SERT +/+ mice. **A)** Illustration of a mouse undergoing the Forced Swim Test, a behavior assay used to assess depressive-like behavior in rodents. **B)** Illustration of a mouse in the Tail Suspension Test, another commonly used assay for evaluating depressive-like behavior. **C)** Box plot comparing immobility times between SERT -/- and SERT +/+ mice in the Forced Swim Test. **D)** Box plot comparing immobility times between SERT -/- and SERT +/+ mice in the Tail Suspension Test.

### Tail Suspension Test

Another measure for depressive behavior was the use of the Tail Suspension Test (TST) (Figure 1B). Mice are suspended by their tails from an aluminum suspension bar, ensuring that they cannot touch the compartment walls or the floor, with their noses positioned 20-25 cm above the floor. Tape, cut to 17 cm, is securely attached to the mouse’s tail, with a 2 cm portion adhering to the tail and the remaining 15 cm used for suspension. White noise generators are used to mask environmental noise. The total duration of immobility and mobility is recorded during behavioral analysis, with mobility defined as active movements aimed at escape, such as trying to reach the walls or strong body shaking. Subtle movements, like pendulum swings or movements limited to the front legs, are not scored as mobility. For each mouse, the observer is blinded to the animal’s group assignment to reduce bias. Statistical analysis was completed in R (R version 4.4.1), where the Mann-Whitney U test was applied using the *wilcox.test* function. A threshold of 0.05 was applied to determine significance. Visualizations were produced using *ggplot2* in R.

### Microbial Sampling and Metagenomic Sequencing

Fecal pellets were collected and stored at −80 °C. For analysis, samples were processed through One Codex using the One Codex Sample Kits to ensure reproducibility and traceability from the lab through to the final results on the One Codex platform. In brief, One Codex extracts DNA, conducts quality control, and sequenced using shotgun sequencing method (with a target depth of 2 million 2x150 bp read pair), enabling species and strain level taxonomic resolution. Sequencing data was analyzed against the One Codex Database, which contains over 115,000 whole microbial reference genomes. A rapid, sensitive k-mer classification algorithm was used to map sequencing reads to reference genomes. To ensure high accuracy, the raw classification results were subjected to rigorous statistical post-processing to filter out false positives caused by contamination or sequencing artifacts. The median read depth was 7,139,050 reads, with a range of 6,023,427 to 8,818,781 reads per sample.

### Diversity Analysis

All analyses of microbial diversity were conducted using QIIME2 (v2024.5)^20^ in an Anaconda environment ^21^. To reduce noise from low-abundance taxa, all taxa that did not meet the frequency cut-off of 0.001 (142,477 reads) were removed. For alpha diversity, rarefaction was performed to ensure even comparison across all samples to a depth of 5,497,439 to retain all samples while ensuring sufficient sequencing coverage. Alpha diversity metrics, including Shannon diversity index, Chao1 species richness index, dominance index, and abundance-based coverage (ACE) index, were calculated using the qiime diversity core-metrics command. Beta diversity metrics, including Bray-Curtis dissimilarity, were also calculated using this core metrics command. After calculating diversity metrics, the results were exported from QIIME2 for statistical analysis in R. Statistical analysis was completed in R (R version 4.4.1), where the Mann-Whitney U test was applied using the *wilcox.test* function for alpha diversity and PERMANOVA in the vegan library was used for beta diversity. A threshold of 0.05 was applied to determine significance. Visualizations of all diversity metrics were produced using *ggplot2* in R.

### Differential Taxonomic Abundance Analysis

For differential abundance analysis, we used MaAsLin2 (Multivariate Association with Linear Models) ^22^, a tool designed for discovering associations between microbial features and metadata variables. We used the species-level feature table and conducted analysis following standard procedures with default parameters, unless otherwise specified. The feature table, metadata, and taxonomy data were imported into R for processing. The data was normalized using centered log-ratio normalization. Associations were based on Cohort (ie. SERT -/- vs. SERT +/+), controlled for sex and litter (fixed_effects = (“Sex”, “Litter”)).

### Microbial Depression Score

A Random Forest classifier using the *caret* package in R was utilized to predict SERT -/- from SERT +/+ mice. The top five differentially abundant microbial species from MaAsLin2 were used for the model. We used a 5-fold cross-validation setup to train the random forest model, optimizing the number of features at each split (mtry parameter) to improve model performance. The final model was evaluated using accuracy scores to assess its predictive performance. The final value used for the model was mtry = 2, with an accuracy of 0.82. The trained random forest model was used to predict the probabilities of samples belonging to the SERT -/- cohort, ranging from 0 to 1, which represented the microbial depression score. We used this score to explore its relationship with behavior outcomes, quantified by the FST score and the TST score. Variable importance, using the mean decrease gini metric, was calculated using the *randomForest* package in R. We used Pearson correlation to assess the strength and direction of the association between the microbial score and each behavioral outcome. We applied the *cor.test()* function in R, specifying method = “pearson” to perform correlation analysis. The correlation coefficients and p-values were reported to summarize the results.

### Bacterial Co-Occurrence Network

To visualize microbial associations and explore differences in microbial interactions between SERT -/- and SERT +/+ mice groups, two bacterial co-occurrence networks were generated. Sparse Co-Occurrence Network Investigation for Compositional data (SCNIC) was used within the QIIME2 environment. Sparse correlations for compositional data (SParCC) was used to calculate relationships between phyla based on abundances from each cohort’s ASV count tables. A correlation threshold of 0.50 was used to retain stronger associations. The network file was imported into Cytoscape for visualization. We used the co-occurrence matrices to construct adjacency matrices, which were padded for compatibility using the numpy library in Python. In order to assess the degree of similarity between the SERT +/+ and SERT -/- co-occurrence matrices, we used the *mantel* function from *skbio.stats.distance* module. We set method=’pearson’ for a Pearson correlation-based Mantel test and specified 999 permutations to assess the statistical significance, returning a correlation coefficient and p-value.

## Results

### SERT -/- mice exhibit depressive-like behaviors

To evaluate the impact of knocking out the SERT gene on behavior in mice, we compared scores for the FST and TST between the experimental cohorts (Figure 1A and Figure 1B), where increased immobility time is indicative of behavioral despair. A significant difference was observed in FST scores (p = 0.004), with SERT -/- mice displaying a higher average score (217.36) compared to SERT +/+ mice (165.89) (Figure 1C). The effect size for the FST scores, measured by Cohen’s d, was 1.75, indicating a large and meaningful difference in depressive-like behaviors between the SERT -/- mice and SERT +/+ controls. In addition to the significant difference observed in the FST, we compared scores for the TST between the two cohorts. While SERT -/- mice had a higher mean TST score (245.27) compared to SERT +/+ mice (214.11), this difference did not reach statistical significance (p = 0.080) (Figure 1D). Despite the lack of statistical significance, the effect size for the TST scores were 0.86, indicating a moderate to large effect between the cohorts. From this analysis, we learned that knocking out the SERT gene in mice leads to differences in depression-like behaviors.

### Alpha diversity, but not beta diversity, is altered by SERT deficiency

To assess the differences in ecological structure of bacterial communities in fecal isolates between SERT -/- and SERT +/+ mice, alpha diversity (within-sample diversity) and beta diversity (between-sample diversity) were measured. Alpha diversity, assessed using abundance-based coverage estimator (ACE) index, Chao1 index, Shannon index, and Simpson Index, revealed that the SERT -/- mice had higher diversity (Figure 2A). SERT -/- and SERT +/+ mice had statistically significant differences in the Shannon index (p = 0.050), while there was no significant difference in the ACE, Chao1, and Simpson indices. The Shannon index captures the number of species present and the evenness of their abundances, so the higher Shannon index value in SERT -/- mice indicates greater microbial diversity. We analyzed beta diversity using Bray-Curtis dissimilarity, which measures the dissimilarity between two communities based on the relative abundance of species (Figure 2B). This beta diversity metric did not reach statistical significance (p= 0.529). Together, these findings suggest that while within-sample (alpha diversity) differences were significant, as measured by the Shannon index, SERT deficiency does not impact the overall beta diversity.

**Figure 2:**
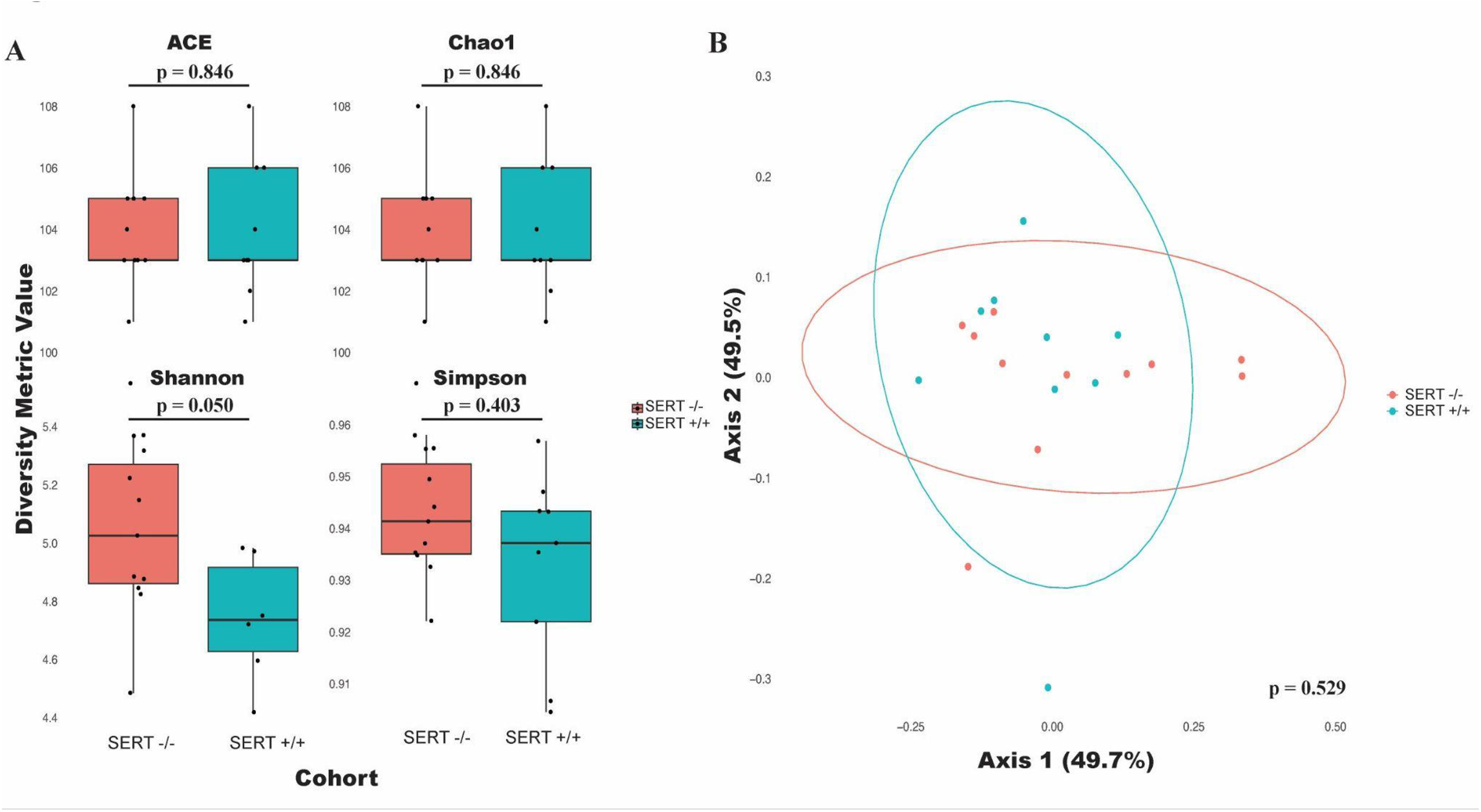
Alpha and beta diversity analysis of gut microbiota in SERT -/- and SERT +/+ mice. **A)** Box plots comparing alpha diversity metrics–ACE, Chao1, Shannon, and Simpson indices–between SERT -/- and SERT +/+ mice. **B)** PCoA plot based on Bray-Curtis dissimilarity showing the clustering of gut microbial communities in SERT -/- and SERT +/+ mice. The ellipses represent 95% confidence intervals for each group.

### Bacterial interaction patterns are altered in SERT deficient mice

Bacterial co-occurrence encompasses the associations between microbial species within the gut environment, forming a network of mutualistic, competitive, or neutral interactions that collectively shape the microbiome’s structure, stability, and functionality. To assess co-occurrence differences, species-species co-occurrence networks were generated for correlated (r > 0.5) bacterial taxa within each cohort (Figure 3A, Figure 3B). The analysis revealed that the SERT +/+ microbiome displayed a more cohesive and interconnected network structure compared to the SERT -/- microbiome, which appeared more fragmented and less interconnected. This suggests that in the SERT +/+ cohort, bacterial species have stronger and more numerous interactions, possibly promoting greater microbial stability and resilience. To quantitatively compare the structure of these co-occurrence networks, Mantel tests were employed to examine the similarity between adjacency matrices derived from each cohort’s co-occurrence network. The Mantel test revealed a statistically significant difference between the cohorts (p = 0.001), indicating a meaningful dissimilarity in the microbial network composition across the groups.

**Figure 3:**
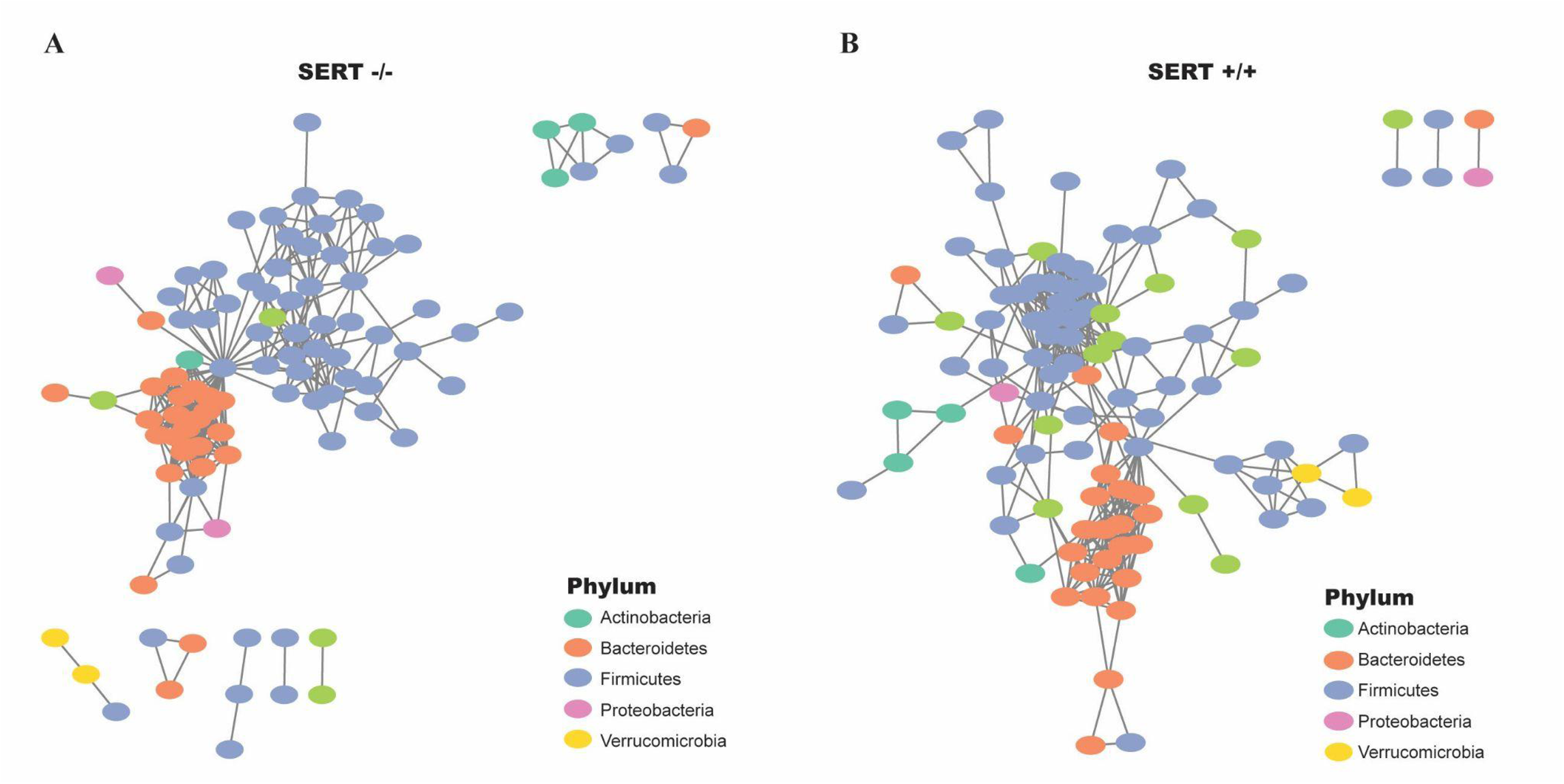
Microbial co-occurence networks in SERT -/- and SERT +/+ mice. **A)** Network plot representing co-occurrence relationships among bacterial taxa in the gut microbiota of SERT -/- mice. Each node represents a bacterial species, colored by phylum, and each edge indicates co-occurrence relationships between species. **B)** Network plot of bacterial co-occurrence relationships in SERT +/+ mice, with nodes and edges structured similarly to panel A.

### SERT deficiency changes the abundance of Firmicutes and Proteobacteria species

The top five phyla were graphed and accounted for greater than 75% of all reads in every sample, exhibiting minor differences in relative abundance between cohorts (Figure 4A). To identify bacterial taxa associated with the SERT -/- phenotype at a more granular level, we conducted differential abundance analysis using MaAsLin2, a statistical framework that assesses multivariable associations between microbial features and metadata while controlling for covariates. MaAsLin2 was applied to species-level abundance data, with SERT -/- versus SERT +/+ cohorts as the primary variable of interest and litter and sex as covariates. Through this analysis two species were differentially abundant, *Clostridium sp. MD294* (p=0.029) and *Acetatifactor MGBC165152* (p=0.048). We used the top 5 abundant bacteria, which included *Clostridium* and *Acetatifactor* based on p-values to generate our microbial depression score. These 5 bacteria were plotted to illustrate their relative abundance between the two groups (Figure 4B). This analysis exhibits some species-level alteration in SERT -/- mice microbial communities.

**Figure 4:**
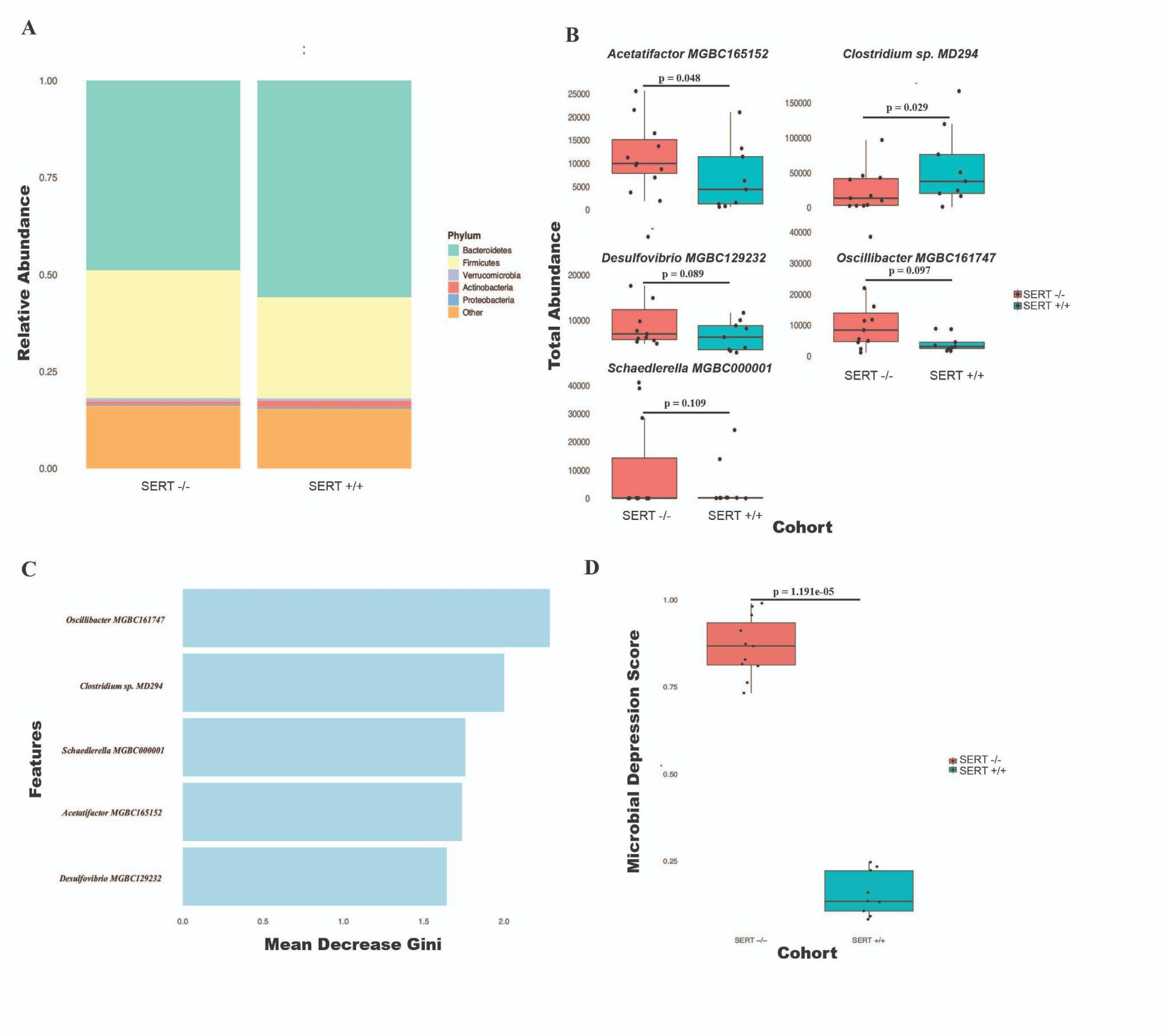
Taxonomic composition differences in SERT -/- and SERT +/+ yields a novel Microbial Depression Score. **A)** Stacked bar plots showing the relative abundance of bacterial phyla in SERT -/- and SERT +/+ mice. **B)** Box plots displaying the total abundance of five bacterial species that differ between SERT -/- and SERT +/+ mice. **C) B**ar chart showing the top five bacterial species ranked by their mean decrease in Gini index, indicating their importance in predicting depressive phenotypes in the random forest model. **D)** Box plot comparing the Microbial Depression Score, derived from the relative abundances of key bacterial species, between SERT -/- and SERT +/+ mice.

### Abundances of five microbial species differentiate SERT -/- mice from SERT +/+ mice

We developed a random forest classifier using the top five differentially abundant bacterial species identified through MaAsLin2 to assess whether microbial composition can accurately differentiate SERT -/- mice from SERT +/+ mice. The selected bacteria were C*lostridium sp.*, *Acetatifactor*, *Desulfovibrio MGBC129232*, *Oscillibacter,* and *Schaedlerella*. The model achieved an accuracy of 0.82 in classifying SERT -/- and SERT +/+ mice, indicating a moderate ability to distinguish between the two groups based on microbiome composition. Then, we derived a microbial depression score, ranging from 0 to 1, using predicted probabilities from the random forest classifier. To evaluate the contribution of each bacterial species to the model’s predictive performance, we calculated the Mean Decrease Gini (MDG) as a measure of variable importance. Clostridium sp. had the highest MDG of 2.284, followed by Schaedlerella (MDG = 1.759), Acetatifactor (MDG = 1.738), Desulfovibrio MGBC129232 (MDG = 1.642), Oscillibacter, and Schaedlerella (Figure 4C). All SERT -/- mice were given microbial depression scores of less than 0.5 with this method (Figure 4D). This analysis demonstrated that these five bacterial species can be used to predict the SERT -/- genotype, highlighting their value as biomarkers of the SERT -/- phenotype.

### Microbial depression score is correlated with depressive phenotypes

We incorporated the calculated microbial depression scores into linear regression models to evaluate its relationship with depressive phenotypes. A Pearson’s correlation analysis revealed a strong positive correlation between the microbial depression score and FST immobility times (Pearson correlation coefficient, r = 0.705), which reached significance and indicated that higher microbial depression scores are associated with increased immobility in the FST (Pearson’s correlation test, p = 0.00052) (Figure 5A). This linear relationship shows that microbial depression score is a strong predictor of FST immobility time. We also examined the association between the microbial depression score and TST immobility times. While a positive correlation was observed (Pearson correlation coefficient, r = 0.401), this trend did not reach statistical significance (Pearson’s correlation test, p = 0.080), suggesting a weaker or less consistent relationship between the microbial depression score and TST performance (Figure 5B). Overall, these results support the microbial depression score as a meaningful tool for assessing microbiome-based depressive traits.

**Figure 5:**
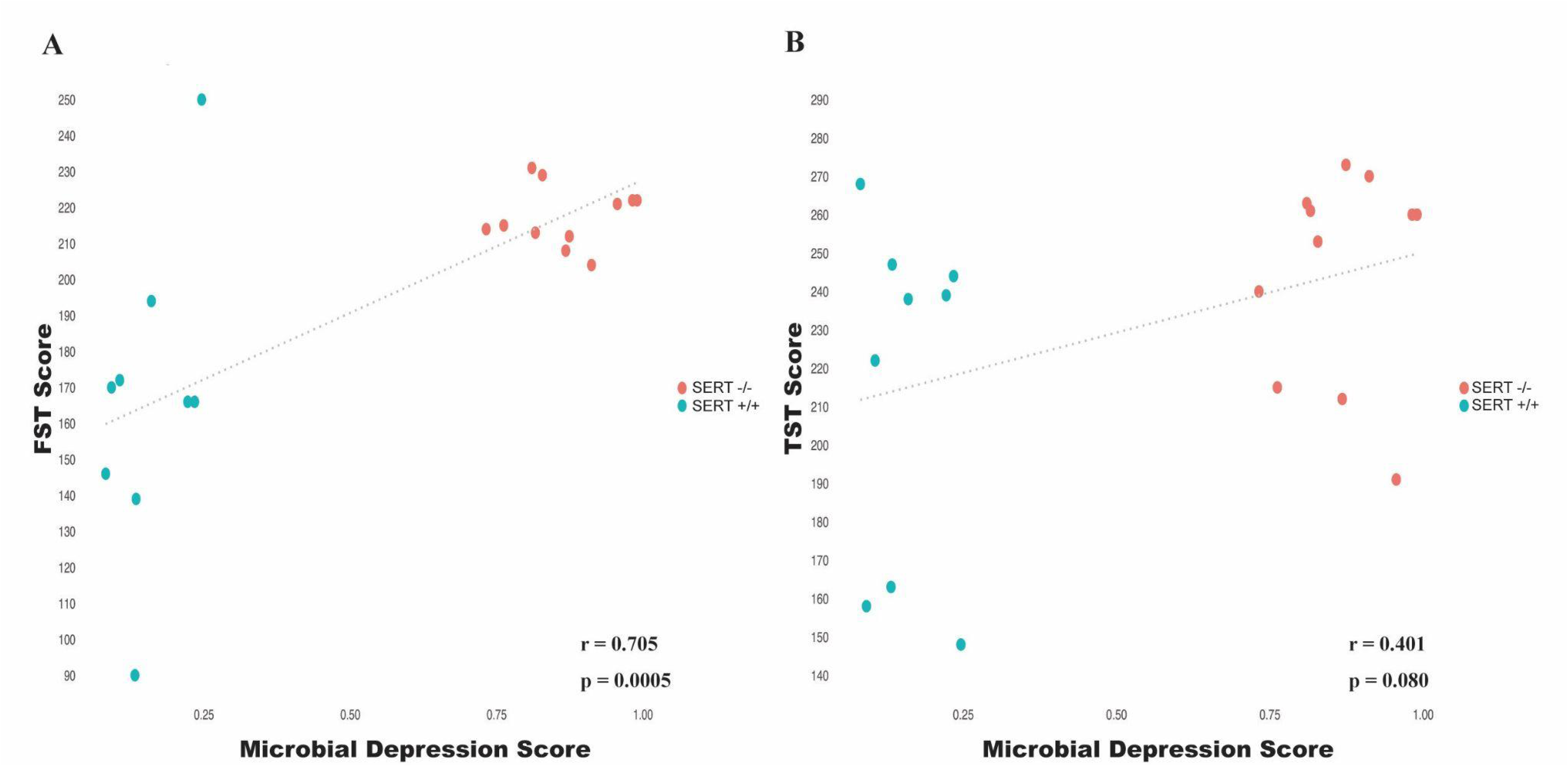
Correlation between Microbial Depression Score and behavioral measures in SERT -/- and SERT +/+ mice. **A)** Scatter plot showing the correlation between the Microbial Depression Score and FST immobility times for SERT -/- and SERT +/+ mice. **B)** Scatter plot showing the correlation between the Microbial Depression Score and TST immobility times for SERT -/- and SERT +/+ mice.

## Discussion

The gut microbiome is increasingly recognized as a critical contributor to the pathogenesis of numerous health disorders. In this study we examined whether genetically-induced depression was associated with microbiome changes, utilizing SERT knockout mice and their litter mate controls. We observed that the loss of the SERT functionality in mice leads to alterations in gut microbial composition, specifically marked by disruptions in the *Firmicutes* and *Proteobacteria* phyla, that was associated with a depressive-like state in SERT -/- mice. By creating a microbial depression score utilizing 5 species, we were able to accurately identify depressive behavioral outcomes in mice.

Behavioral analyses confirmed that SERT -/- mice exhibited depressive-like behaviors, as demonstrated by significantly higher immobility times in both the FST and TST compared to SERT+/+ controls. These results align with prior findings that SERT deficiency disrupts serotonin regulation, promoting behavioral despair^9^. The larger effect size observed in the FST suggests this test may more sensitively capture depressive behavior in SERT-/- mice, while smaller, non-significant differences in the TST highlight the need for multifaceted behavioral assessments. These findings support the utility of SERT-/- mice as a model for investigating the biological mechanisms underlying depression.

The depressive-like symptoms observed may stem from alterations in their gut microbial community that is driven by SERT deficiency. Without functional SERT, serotonin reuptake from the gut lumen is impaired, leading to persistently elevated extracellular serotonin levels. This increase in serotonin can create a selective environment that promotes the growth of specific microbial species adapted to serotonin-rich conditions. One study found that serotonin affects bacterial growth in-vitro in a concentration-dependent and species-specific manner ^23^. To compound these effects, research has shown that serotonin is vital to promoting immune system development in early life through T cell immune tolerance to commensal bacteria and dietary antigens ^24^. This immune tolerance is essential for preventing unnecessary inflammatory responses to non-pathogenic microbes. Therefore, serotonin dysregulation not only modifies gut microbial communities but also alters the microbiota’s interactions with host immune cells, contributing to depressive phenotypes through brain-gut interactions.

Our microbial analysis revealed significant differences in bacterial co-occurrence and alpha diversity between the two genotypes. While beta diversity did not differ significantly between SERT-/- and SERT+/+ mice, distinct differences in bacterial co-occurrence patterns indicate how the structure and stability of the gut microbiome may be affected by SERT deficiency. From this analysis, we observed distinct structural patterns in bacterial interactions within each cohort, highlighting ecological relationships among bacterial taxa that may reflect shifts in microbiome stability and functionality. The greater interconnectedness observed in SERT +/+ mice suggests a more stable and resilient microbial community, while the fragmented network in SERT -/- mice may reflect a dysbiotic state that could contribute to the altered gut-brain interactions observed in this group. Studies on alpha diversity changes in depression show mixed results in both mice and humans ^25,26^. However, in our study, we observed increased alpha diversity in SERT -/- mice. Higher microbial diversity is typically associated with a resilient and stable gut ecosystem; however, in this case, decreased serotonin in SERT -/- mice may promote a dysbiotic shift toward potentially pathogenic or pro-inflammatory bacteria. We observed such a shift towards pro-inflammatory microbial phyla, where our data revealed an increase in *Firmicutes* in SERT-/- mice, a phylum that previous studies have associated with depression and metabolic disturbances. Indeed, previous studies have shown that the abundance of Firmicutes is the most important phylum that is correlated with depression, where taxonomic biomarkers comparing healthy and depressed people were all members of the *Firmicutes* phyla ^27^. These shifts toward pro-inflammatory microbiota lead to depression through numerous mechanisms. The release of pro-inflammatory cytokines into the bloodstream can cross the blood-brain barrier and influence the CNS ^28^. In addition, inflammatory cytokines can activate indoleamine 2,3-dioxygenease (IDO), which diverts tryptophan away from serotonin synthesis and toward the kynurenine pathway, which is increased in patients with depression ^29^.

One of our study’s key findings is the development of a microbial depression score, derived from the abundances of *Acetatifactor MGBC165152, Clostridium sp. MD294, Desulfovibrio MGBC129232, Oscillibacter MGBC161747, and Schaedlerella MGBC000001*, which positively correlated with FST and TST immobility times. We observed that these species were associated with major changes in depressive behavior outcomes. This highlights the potential utility of microbiome-based markers in depressive behavior. The most differentially abundant species between SERT-/- and SERT+/+ mice were *Acetatifactor MGBC165152* and *Clostridium sp. MD294,* both belonging to *Firmicutes* phylum*. Acetatifactor MGBC165152,* a gram-positive member of the *Firmicutes* phylum, was significantly increased in SERT -/- mice. *Acetatifactor* has been implicated in promoting dextran sulfate sodium (DSS)-induced colitis, which damages the colon-lining and exacerbates inflammation ^30^. *Clostridium sp. MD294,* a gram-positive member of the *Firmicutes* phylum, was the only species of the five used for the microbial depression score with decreased abundance in SERT-/- mice, aligning with findings that reduced levels of certain *Clostriudium* species correlate with depression ^31^.

Furthermore, our microbial depression score included abundances for the *Desulfovibrio MGBC129232* species, a member of the Proteobacteria phylum. The genus *Desulfovibrio* is typically regarded as a harmful genus due to its association with inflammatory diseases, such as obesity ^32^, and Ulcerative Colitis ^33^. *Desulfovibrio* utilizes short-chain fatty acids, sulfur-containing amino acids, and inorganic sulfates as electron donors to produce hydrogen sulfide (H₂S) ^34^. This metabolite has been shown to promote inflammation by activating pathways such as ERK1/2, p38 MAPK, and ERK-NF-kB, leading to the synthesis of pro-inflammatory cytokines such as TNF-α, IL-1β, and IL-6 ^35^. Additionally, as a gram-negative bacterium, *Desulfovibrio* contains Lipopolysaccharide (LPS) in its outer membranes, which induces inflammatory responses in host cells ^36,37^. *Schaedlerella MGBC000001*, also within the *Firmicutes* phylum and a member of the *Lachnospiraceae* family, was incorporated in the microbial depression score. While research on *Schaedlerella* itself is limited, related taxa within *Lachnospiraceae* (e.g., *Anaerostipes, Blautia, Dore*a) have been positively associated with major depressive disorder ^38^, suggesting that *Schaedlerella* could play a similar role in depression. *Oscillibacter MGBC161747*, another gram-negative species in the *Firmicutes* phylum, had the greatest feature importance in calculating the microbial depression score. Prior studies indicate that *Oscillibacter* are positively correlated with increased gut permeability ^39^, allowing bacterial components and metabolites to cross the gut barrier, resulting in “leaky gut.” This phenomenon is linked to low-grade inflammation and neuroinflammatory responses that exacerbate depressive symptoms.

While our study presents compelling evidence linking gut microbiota to depressive phenotypes in SERT -/- mice, there are some limitations. To truly prove causality, future studies could implement a fecal microbiota transplant, transferring the microbiota from SERT -/- mice into SERT +/+ wildtype mice. This approach would clarify whether the altered gut microbiome composition directly drives the behavioral changes observed in SERT -/- mice. Additionally, the inclusion of a validation set of mice to test our microbial depression score on would add to the robustness of the score. Furthermore, our study focused on the gut microbiota, but did not account for potential interactions between the gut and other physiological systems, such as the immune or endocrine systems, which could also influence depression-like behaviors.

A previous study also demonstrated that SERT deficiency alters the gut microbiome, revealing shifts in bacterial abundance and community structure; however, that study focused on the links between microbial changes and metabolic syndrome ^40^. Additionally, the previous study co-housed mice by genotype, mixing littermates from differ dams, potentially introducing cage effects that may have influenced observed differences. In contrast, our study housed mice by litter and genotype, reducing environmental confounds and allowing us to capture more biologically significant differences associated with SERT deletion. We also introduced behavior assays to see how the microbial shifts that both our study and their study saw can be associated with a depressive phenotype.

Collectively, our study provides evidence that gut microbiota are key modulators of depressive behaviors in SERT-/- mice. This is clinically relevant because selective serotonin reuptake inhibitors (SSRIs) are a common prescription to treat depression, which target SERT in order to increase the amount of serotonin available in the synaptic cleft. However, we see that this may lead to gut microbiota alterations that may lead to exacerbated depressive behaviors, potentially explaining varying clinical responses to SSRI. Future research may investigate the mechanistic underpinnings of these microbial effects, particularly their influence on serotonin metabolism and signaling pathways, or transferring these bacteria to germ-free mice via oral gavage to determine whether they induce depressive-like behaviors. Additionally, expansion of this research to human cohorts will be crucial for validating the relevance of these microbial signatures in clinical contexts. A deeper understanding of these interactions could pave the way for microbiome-targeted therapies as innovative approaches to treating depression.

## Data Availability

Sequencing data will be deposited to publicly available repository upon publication

## Notes

### Competing Interest Statement

The authors have declared no competing interest.

